# Dynamics of circulating tumor cell subsets defined by PSMA and EpCAM predict survival in metastatic castration-resistant prostate cancer patients treated with ^177^Lu-PSMA-617

**DOI:** 10.64898/2026.09.14.751487

**Authors:** Madeline J. Lee, Lauren Yu, Nicholas A. Zorko, Scott Dehm, Justin H. Hwang, Justin M. Drake, Emmanuel S. Antonarakis, Ali T. Arafa

**Affiliations:** Division of Hematology, Oncology and Transplantation, Department of Medicine, University of Minnesota, Minneapolis, MN 55455, USA; Masonic Cancer Center, University of Minnesota, Minneapolis, MN 55455, USA; Department of Laboratory Medicine and Pathology, University of Minnesota, Minneapolis, MN 55455, USA; Department of Urology, University of Minnesota, Minneapolis, MN 55455, USA; Astrin Biosciences, Saint Paul, MN 55114, USA; Department of Pharmacology, University of Minnesota, Minneapolis, MN 55455, USA

**Keywords:** Radioligand therapy, mCRPC, holographic image analysis

## Abstract

**Background:** ^177^Lu-PSMA-617 radioligand treatment represents a major advancement in the management of metastatic castration-resistant prostate cancer (mCRPC), but key questions remain about tumor-cell dynamics and evolution in patients who have received this drug. Here, we used a circulating tumor cell (CTC) capture assay to assess the prognostic capacity of CTC subset dynamics following ^177^Lu-PSMA-617 treatment in a cohort of mCRPC patients.

**Methods:** Using an AI-empowered holographic imaging platform combined with in-flow protein marker analysis, we prospectively analyzed serial CTC samples from patients receiving ^177^Lu-PSMA-617 therapy at pre-treatment and on-treatment time points (n=39 of 100 enrolled patients). PSMA and EpCAM proteins in CTCs were determined by immunofluorescence staining. Proportions and absolute counts of CTC subsets delineated by PSMA and EpCAM proteins were assessed in PSA50 responders (n=19) and non-responders (n=20). Within-patient changes in CTC subsets were compared using Wilcoxon signed-rank tests with Benjamini-Hochberg correction. Associations of CTC subset abundance and dynamics with overall survival were assessed using Kaplan-Meier estimates with log-rank tests and univariable Cox proportional hazards models.

**Results:** PSMA and EpCAM stratified CTCs into subsets, some of which were associated with survival outcomes following ^177^Lu-PSMA-617. We observed an increase in the proportion of PSMA−/EpCAM+ CTCs following ^177^Lu-PSMA-617, with a larger increase occurring in PSA50 responders (6.2% to 18.7%; p=0.041) than non-responders (7.9% to 15.4%; p=0.023). This corresponded to a decrease in the proportion of PSMA+/EpCAM-CTCs, whereas the proportion of PSMA+/EpCAM+ CTCs did not decline. On-treatment counts of >5 CTCs, relative to ≤5 CTCs, were associated with shorter overall survival in the PSMA+/EpCAM+ (HR 3.56, 95%CI 1.32-9.60, p=0.012) and PSMA+/EpCAM-(HR 7.50, 95%CI 0.96-52.99, p=0.054) subsets. Finally, on-treatment declines in PSMA+/EpCAM+ (HR 0.74, p=0.49) and PSMA+/EpCAM-(HR 0.61, p=0.215) CTC subsets were not significantly associated with overall survival relative to sustained high levels of these cells.

**Conclusions:** In mCRPC patients receiving ^177^Lu-PSMA-617, dynamic changes in CTC subsets delineated by PSMA and EpCAM expression may have prognostic value, justifying additional evaluation.

## Introduction

The advent of ^177^Lu-PSMA-617 therapy represents a major advancement in the management of metastatic castration-resistant prostate cancer (mCRPC) and has more recently been approved in the United States for metastatic hormone-sensitive prostate cancer (mHSPC). ^177^Lu-PSMA-617, a radioligand therapy which delivers lutetium-177 to PSMA-expressing tumor cells, has been shown to extend both progression-free and overall survival in the mCRPC state^1–3^ as well as progression-free survival in the mHSPC state^4,5^. However, the effects of ^177^Lu-PSMA-617 on the landscape of PSMA expression on tumor cells and the prognostic value of these changes remain largely unexplored.

A variety of non-invasive methods can be used to characterize tumor populations and/or function as prognostic indicators, including analysis of circulating tumor DNA (ctDNA)^6,7^ and plasma-based extracellular vesicles (EVs)^8,9^. Post hoc analysis of the TheraP trial (a randomized trial of ^177^Lu-PSMA-617 versus cabazitaxel) found that ctDNA fraction at baseline was negatively correlated with PSA50 response rates as well as progression-free survival (PFS), and identified specific genetic mutations that were significantly associated with PFS and overall survival (OS)^10^. Previous work conducted by our group assessed the ability of EV-derived proteins to predict outcomes to ^177^Lu-PSMA-617 therapy, resulting in the identification of a suite of cell-surface molecules (including PSMA, B7-H3, Trop-2, and STEAP1) whose expression was predictive of worse OS^8^. While these methods provide prognostic value as well as valuable insights into changes induced by ^177^Lu-PSMA-617 therapy, they do not directly examine PSMA expression on tumor cells, nor do they assess how the proportion of PSMA-positive tumor cells may change over time when exposed to a PSMA-targeting therapy.

Circulating tumor cells (CTCs) provide another avenue to study metastatic malignancies without the need for invasive interventions such as biopsies and offer unique advantages over analysis of ctDNA and plasma-based EVs. Most notably, CTCs present the opportunity to directly assess tumor cells at the single-cell level^11^. This allows for analysis such as quantification of single or co-expressed proteins^12^, transcriptomic profiling^13,14^, and single-cell mutational analysis^15,16^. Thus far, studies of CTCs in mCRPC have largely focused on the associations between total CTC count and OS, finding that higher CTC counts are significantly predictive of worse OS^17–19^. While the value of total baseline CTC enumeration as a prognostic indicator has been demonstrated in many systemic-therapy settings, few studies have examined the prognostic utility of changes in PSMA expression on CTCs in the context of ^177^Lu-PSMA-617 treatment.

Here, we conducted a prospective correlative biomarker study in which CTCs were collected at baseline and on-treatment time points in a cohort of 100 mCRPC patients receiving ^177^Lu-PSMA-617, of which 39 patients had paired CTC sample analyses. To identify CTCs, we employed a method that applies a neural network to holographic and immunofluorescent images of erythrocyte-depleted blood samples, resulting in the detection of CTCs with high fidelity while retaining heterogeneity^20^. We identify subpopulations of CTCs defined by expression of PSMA and EpCAM, and we explore the associations between the abundances of these subsets and overall survival outcomes as well as PSA50 response rates. These data illustrate the impact of ^177^Lu-PSMA-617 treatment on CTC subsets delineated by PSMA and EpCAM expression over time, and demonstrate the potential prognostic value in analyzing subpopulations of CTCs in this setting.

## Materials and Methods

### Study participants

We conducted a prospective study of 100 mCRPC patients receiving 177Lu-PSMA-617 (University of Minnesota IRB number: Study00013815). All participants provided written informed consent, and the study was conducted in accordance with the Declaration of Helsinki. Patients could elect to provide a second blood collection 12-18 weeks after treatment initiation. A total of 39 patients provided paired baseline and on-treatment samples. Treatment response was classified according to PSA50 response criteria (per PCWG3), defined as a ≥50% decline in serum PSA from baseline^21^. Overall survival (OS) was defined as the time from treatment initiation to death of any cause. Patients who were alive at the time of data cutoff were censored at last follow up.

### Survival analysis

Survival analysis was performed using the *survminer* and *survival* R packages. Hazard ratios (HR) and 95% confidence intervals (95%CI) were estimated by fitting Cox proportional hazards regression models to the data. Comparisons between Kaplan-Meier estimates were performed using the log-rank test. Survival models were univariable; no adjustment was made for baseline prognostic covariates, and p values from the survival analyses were not corrected for multiple comparisons. Given the sample size and the number of comparisons performed, all survival results should be regarded as exploratory.

### CTC isolation

Peripheral blood samples (7.5 mL per draw) were processed using a previously described circulating tumor cell enrichment and holographic imaging workflow (Astrin Biosciences, Saint Paul, MN)^20^. Following microfluidic enrichment, CTC candidates were identified by holographic imaging and subsequently stained with fluorescently conjugated antibodies targeting prostate specific membrane antigen (PSMA) and epithelial cell adhesion molecule (EpCAM). CTCs were classified according to PSMA and EpCAM expression into three phenotypic populations: PSMA⁺/EpCAM⁺, PSMA⁺/EpCAM⁻, and PSMA⁻/EpCAM⁺. The abundance of each phenotype was quantified per 7.5 mL of whole blood for each patient and used for subsequent analyses.

### Stratification by CTC subset abundance

For each of the three CTC subsets analyzed (PSMA+ EpCAM+, PSMA+ EpCAM-, and PSMA-EpCAM+), patient samples from the pre- and on-treatment time points were stratified into two groups based on the number of CTCs recovered in 7.5 mL blood: >5 CTCs (“High”) or ≤5 CTCs (“Low”). This CTC count threshold was selected because it has been previously described as being clinically significant in the setting of mCRPC^17^. Patients were then further stratified into 4 total groups based on the dynamics of their CTC counts in their pre- and on-treatment samples. For each CTC subset, patients were classified as either Low-Low (≤5 CTCs pre-treatment, ≤5 CTCs on-treatment); Low-High (≤5 CTCs pre-treatment, >5 CTCs on-treatment); High-Low (>5 CTCs pre-treatment, ≤5 CTCs on-treatment); or High-High (>5 CTCs pre-treatment, >5 CTCs on-treatment).

### Statistical analysis

Comparisons of CTC subset proportions and counts between the pre- and on-treatment time points were made within patients and were performed using the Wilcoxon signed-rank test, with Benjamini-Hochberg correction applied across comparisons. Comparisons between PSA50 responders and non-responders were performed using the Wilcoxon rank-sum test. All statistical analyses were performed in R, and two-sided p values <0.05 were considered statistically significant.

## Results

### Clinical characteristics

A total of 100 patients with mCRPC receiving ^177^Lu-PSMA-617 were enrolled, of whom 39 provided paired pre-treatment and on-treatment blood samples and constituted the analytic cohort for this study. Baseline clinical and pathologic characteristics of these 39 patients are summarized in Supplemental Table 1. The median age at study enrollment was 73 years (range 50-94), the median baseline serum PSA was 78 ng/mL (range 0.00-5,000), and the median baseline alkaline phosphatase was 123 U/L (range 41-1,920). Most patients had Gleason grade group 4 or 5 disease at diagnosis (28/39, 72%) and an ECOG performance status of 0 or 1 (27/39, 69%). 19 patients (49%) achieved a PSA50 response and 20 (51%) did not.

### ^177^Lu-PSMA-617 treatment and CTC composition

We analyzed the composition of CTCs (and their changes over time) separately in PSA50 responders (n=19) and non-responders (n=20) using both the baseline and on-treatment time points. Our analysis identified three major subpopulations of CTCs based on PSMA and EpCAM proteins: PSMA+/EpCAM+ cells, PSMA+/EpCAM-cells, and PSMA−/EpCAM+ cells (Fig. 1A-D). At baseline, prior to ^177^Lu-PSMA-617 treatment, PSMA+ cells comprised >90% of CTCs on average, with only one patient having fewer than 75% PSMA+ cells (Fig. 1A, C). Patients averaged approximately 40% EpCAM+ cells, though this was much more variable and ranged between 0-100%. Among PSA50 responders, 16/19 patients had detectable CTCs from all three subsets at baseline, while 2/19 had only PSMA+/EpCAM+ and PSMA+/EpCAM-CTCs, and 1/19 had only PSMA+/EpCAM-CTCs (Fig. 1A). At the second time point, 14/19 responders had all three subsets represented, 3/19 had only PSMA+/EpCAM+ and PSMA+/EpCAM-CTCs, and 2/19 had no detectable CTCs (Fig. 1B). Within the PSA50 non-responder population, a lower proportion of patients (13/20) had CTCs from all three subsets in their pre-treatment sample, while 4/20 patients had only PSMA+/EpCAM+ and PSMA+/EpCAM-CTCs, 2/20 had exclusively PSMA+/EpCAM-CTCs, and one patient had no detectable CTCs at baseline (Fig. 1C). At the second time point, 18/20 non-responders had CTCs from all three subsets recovered, while one patient had only PSMA+/EpCAM+ and PSMA+/EpCAM-CTCs and one patient had only PSMA+/EpCAM-CTCs (Fig. 1D).

**Figure 1:**
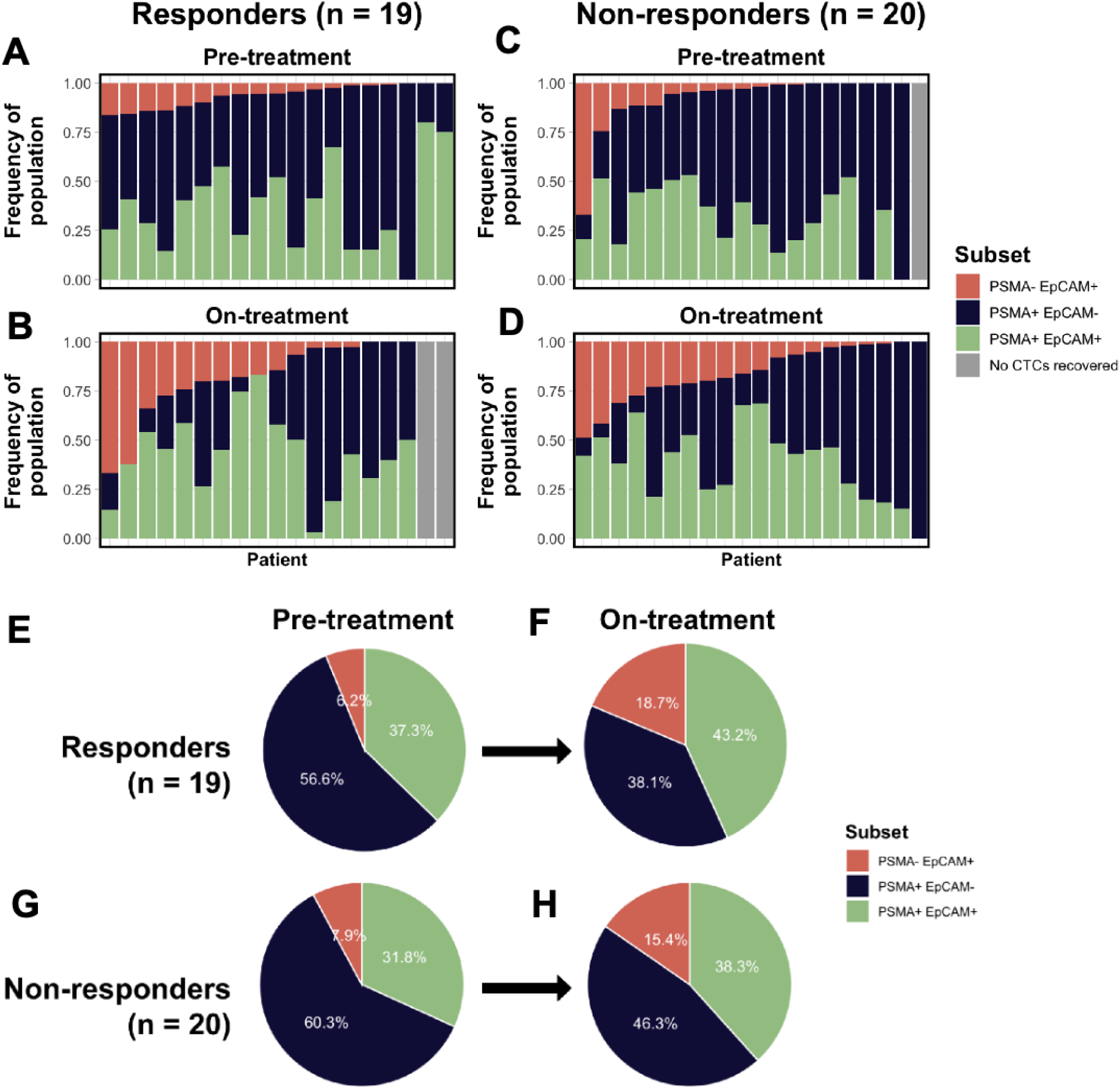
^177^Lu-PSMA-617 treatment alters CTC composition. A-D: Stacked bar plots showing the frequencies of 3 CTC subsets (PSMA+ EpCAM+; PSMA+ EpCAM-; PSMA-EpCAM+) in PSA50 responders (A-B) and non-responders (C-D) at pre- and on-treatment time points. Data are presented as fraction of total CTCs represented by each cell subset. Gray bars represent patients for whom no CTCs of a given subset were recovered. E-H) Pie charts showing the evolving distribution of CTC subsets in PSA50 responders (E-F) and non-responders (G-H) at pre- and on-treatment time points.

Following ^177^Lu-PSMA-617 treatment, the mean proportion of PSMA−/EpCAM+ CTCs increased approximately three-fold, from 6.2% in pre-treatment samples to 18.7% in on-treatment samples, in PSA50 responders (Fig. 1E-F, Fig. 2A), while the same population expanded from a mean of 7.9% pre-treatment to 15.4% on-treatment in non-responders (Fig. 1G-H, Fig. 2B). This increase was statistically significant in both PSA50 responders (p=0.041; Fig. 2A) and non-responders (p=0.023; Fig. 2B). The magnitude of change in the proportion of this PSMA-population was larger in responders relative to non-responders, although the difference between responders and non-responders did not reach statistical significance (p=0.277).

**Figure 2:**
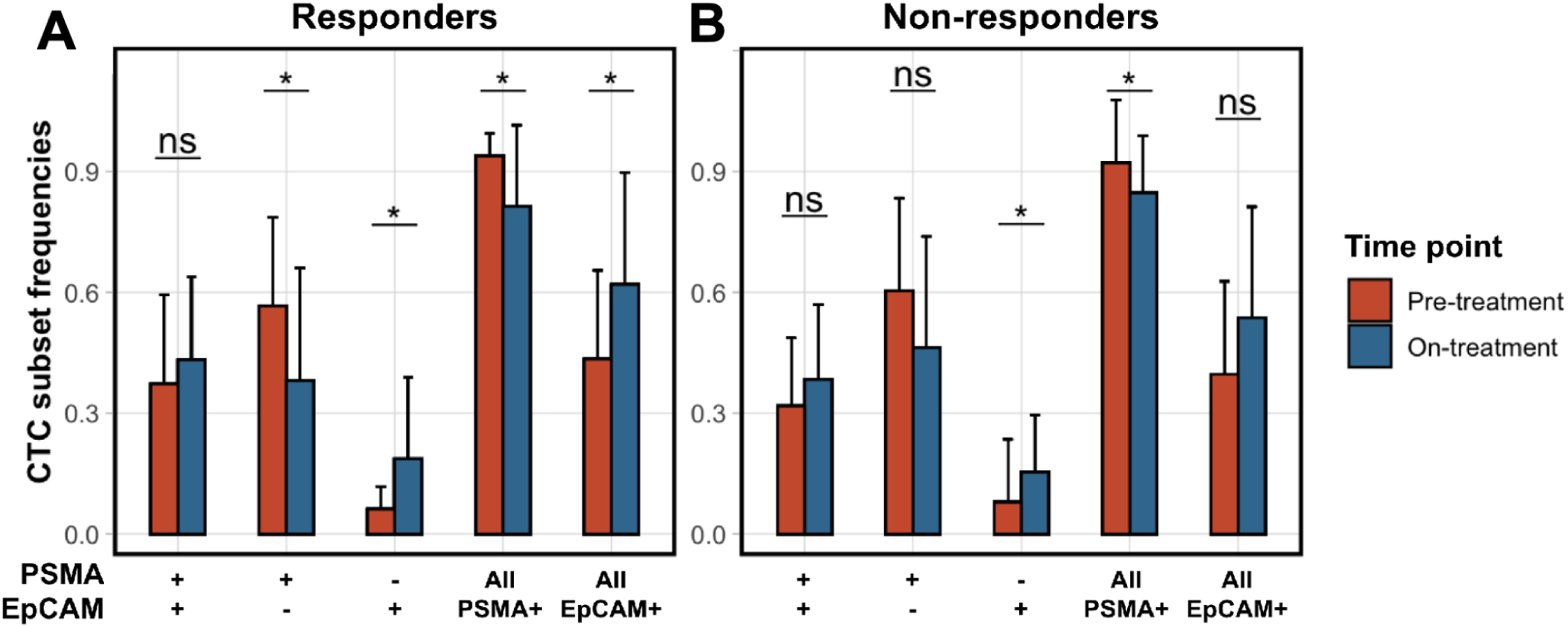
Frequency of PSMA-EpCAM+ CTCs increases during ^177^Lu-PSMA-617 treatment. Bar plots showing the frequencies of CTC subsets defined by surface expression of PSMA and EpCAM in pre- and on-treatment samples for PSAS0 responders (A; n=19) and non-responders (B; n=20). Error bars represent one standard deviation. Significance values were determined using a Wilcoxon rank-sum test with the Benjamini-Hochberg correction.

Concurrent with the observed increase in PSMA−/EpCAM+ CTC proportion was a decrease in the fraction of PSMA+/EpCAM-CTCs in on-treatment samples. The average proportion of PSMA+/EpCAM-CTCs in the baseline pre-treatment sample was 56.6% in PSA50 responders and 60.3% in non-responders; this decreased to 38.1% among responders (p=0.039) and 46.3% among non-responders (p=0.106) at the on-treatment time point (Fig. 1E-H; Fig. 2). Meanwhile, the proportion of PSMA+/EpCAM+ cells increased slightly from 37.3% to 43.2% in PSA50 responders (p=0.326) and from 31.8% to 38.3% in non-responders (p=0.361) (Fig. 1E-H, Fig. 2).

### PSMA+ but not PSMA-CTCs are less abundant following ^177^Lu-PSMA-617 treatment

We next assessed the change in absolute CTC numbers for each cell subset between pre- and on-treatment samples (Fig. 3). Consistent with ^177^Lu-PSMA-617’s mechanism of action^22,23^, the majority of patients experienced a decrease in PSMA+/EpCAM+ or PSMA+/EpCAM-CTC counts following radioligand therapy (23/39 and 27/39 patients, respectively) (Fig. 3A-B). A greater fraction of PSA50 responders (15/19) experienced a decrease in the number of total PSMA+ cells compared to PSA50 non-responders (13/20) (Fig. 3C). Conversely, only 11/39 patients exhibited a numerical decrease in PSMA−/EpCAM+ CTCs (Fig. 3D). The decrease in PSMA-CTCs was similar in both PSA50 responders (5/19) and non-responders (6/20), suggesting that the presence of a quantitative decrease in PSMA-CTCs may not be associated with efficacy of ^177^Lu-PSMA-617.

**Figure 3:**
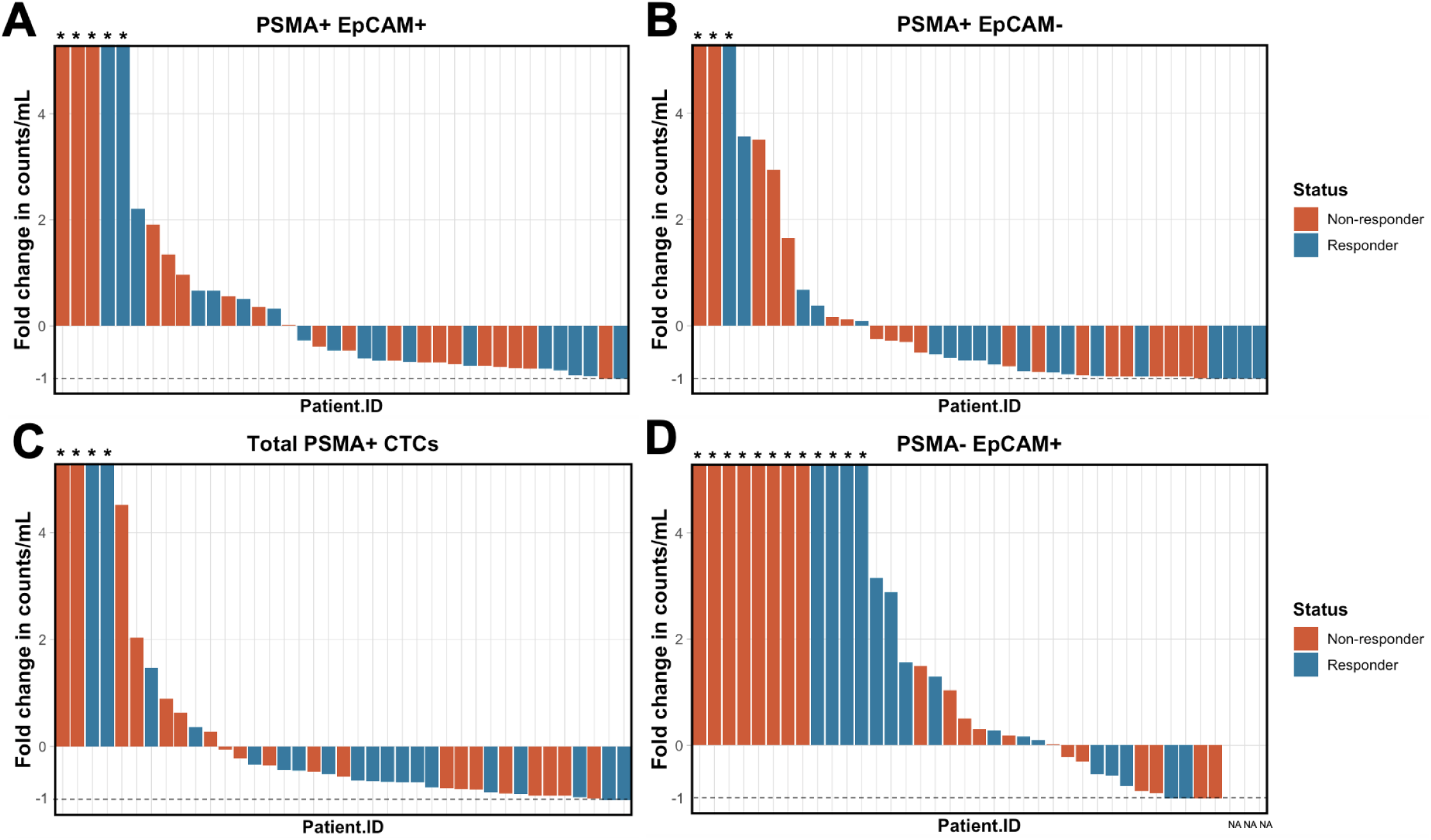
PSMA-but not PSMA+ CTCs are more abundant following ^177^Lu-PSMA-617 treatment. Waterfall plots showing the fold change in counts per ml of three CTC subsets: A) PSMA+ EpCAM+; B) PSMA+ EpCAM-; C) PSMA-EpCAM+. Bars are colored by PSA50 response status. Asterisks indicate cases where the fold increase was greater than 5. NAs indicate cases where zero CTCs were recovered from that subset in both the pre- and on-treatment samples. Dashed line indicates the threshold for a 100% decrease.

**Supplemental Figure 1:**
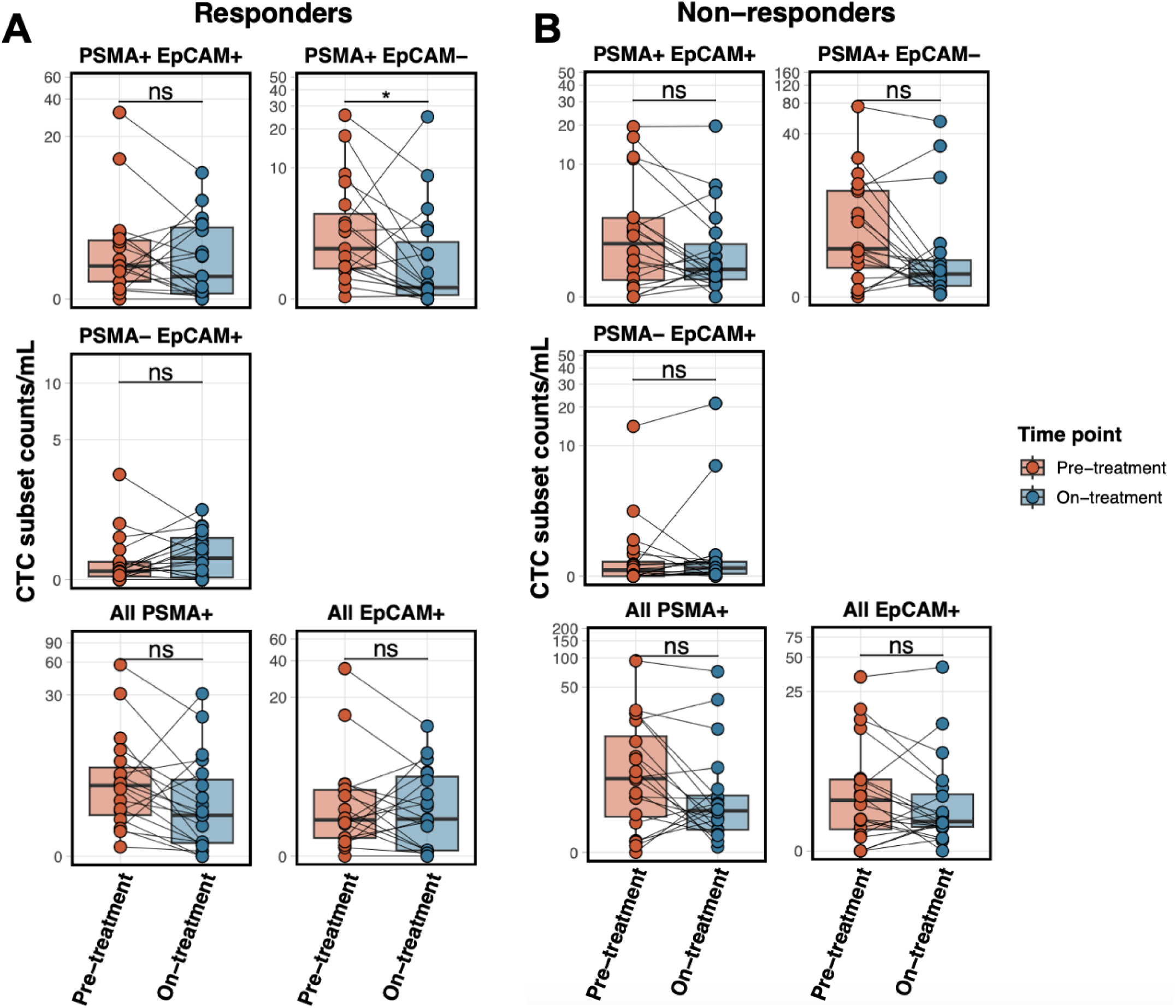
PSMA+ CTCs remain following ^177^Lu-PSMA-617 therapy. Boxplots showing the counts per ml recovered from CTC subsets defined by PSMA and EpCAM expression in PSA50 responders (A; n=19) and non-responders (B; n=20). Lines connect pre- and on-treatment samples from individual patients. Significance values were determined using a Wilcoxon rank-sum test with the Benjamini-Hochberg correction. Data are shown on a pseudolog scale to account for heteroscedasticity.

### PSMA+ CTCs persist following ^177^Lu-PSMA-617 therapy

A key outstanding question is whether PSMA remains a targetable tumor antigen following radioligand treatment in patients who respond to ^177^Lu-PSMA-617 therapy. In this regard, we found that, although 15/19 PSA50 responders experienced a decrease in the number of PSMA+ CTCs in their on-treatment sample relative to pre-treatment (p=0.118), only two of these patients had zero recoverable PSMA+ CTCs at the second time point (Fig. S1A). Similarly, 13/20 PSA50 non-responders experienced a decrease in PSMA+ CTC counts (p=0.149), but none of these patients had zero recoverable PSMA+ CTCs at the second time point (Fig. S1B). Overall, these data suggest that while ^177^Lu-PSMA-617 treatment decreases the number of PSMA+ CTCs and results in a compensatory increase of PSMA-CTCs, there are still persistent PSMA+ tumor cells available for targeting by further lines of PSMA-directed therapy.

### Patients with ≤5 PSMA+ CTCs following ^177^Lu-PSMA-617 treatment have more favorable survival outcomes

We next explored the effects of CTC subset count on overall survival by stratifying patients into those who had >5 CTCs from a given subgroup recovered in their on-treatment sample versus those with ≤5 CTCs in their second sample (Fig. 4). This cutoff was selected based on work from de Bono et al. demonstrating 5 CTCs per 7.5 mL blood to be a clinically significant threshold in mCRPC for prognostication^17^. To this end, we found that patients with >5 CTCs on-treatment within any subset had worse overall survival, although the effect size was larger in the PSMA+/EpCAM+ population (HR 3.56, 95%CI 1.32-9.60, p=0.012; Fig. 4A) and the PSMA+/EpCAM-population (HR 7.50, 95%CI 0.96-52.99, p=0.054; Fig. 4B) than in the PSMA-subset (HR 1.89, 95%CI 0.79-4.55, p=0.154; Fig. 4C); only the PSMA+/EpCAM+ comparison reached statistical significance, and the wide confidence interval for the PSMA+/EpCAM-estimate reflects the small number of patients and events in that stratum. This may reflect the increased tumor burden of patients with higher CTC counts and the inability of ^177^Lu-PSMA-617 to sufficiently clear CTC populations below the 5 CTC threshold.

**Figure 4:**
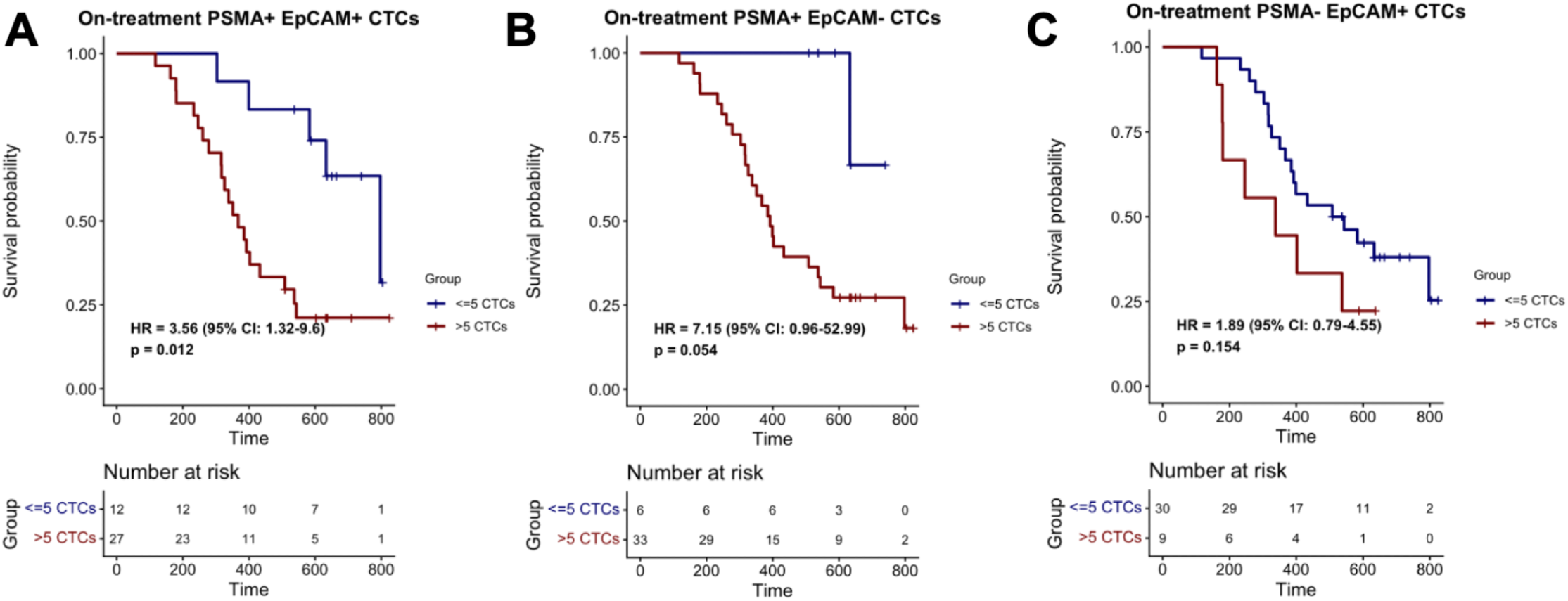
Patients with <=5 PSMA+ CTCs following ^177^Lu-PSMA-617 treatment have better survival outcomes. Kaplan-Meier survival curves showing overall survival based on stratification by number of CTCs recovered in the on-treatment sample (<= 5 or >5). Data are shown for PSMA+ EpCAM+ CTCs (A), PSMA+ EpCAM-CTCs (B), and PSMA-EpCAM+ CTCs (C). Comparisons are made using the log-rank test.

### Changes in CTC subsets between pre- and on-treatment samples are associated with survival outcomes

Finally, we stratified patients into four groups based on changes in CTC counts between pre- and on-treatment samples. Using the cutoff of >5 CTCs, patients were categorized as High-High (>5 CTCs pre-treatment, >5 CTCs on-treatment); Low-Low (≤5 CTCs in both pre- and on-treatment samples); Low-High (≤5 CTCs pre-treatment, >5 CTCs on-treatment); and High-Low (>5 CTCs pre-treatment, ≤5 CTCs on-treatment). We then performed survival analyses to assess whether changes in the abundance of each CTC subset correlated with overall survival.

In both PSMA+ subsets, having ≤5 CTCs pre-treatment was associated with better overall survival (Fig. 5A-B). In the PSMA+ EpCAM+ population, there was a small additional survival benefit in the Low-Low group (HR=0.18, 95%CI 0.02-0.71, p=0.01) versus the Low-High group (HR=0.37, 95%CI 0.11-1.00, p=0.05). Patients in the High-Low groups had a slight but nonsignificant survival benefit relative to the High-High group (PSMA+/EpCAM+: HR=0.74, 95%CI 0.27-1.76, p=0.49; PSMA+/EpCAM-: HR=0.61, 95%CI 0.27-1.33, p=0.215). These exploratory data raise the possibility that pre-treatment PSMA+ CTC count carries prognostic information beyond that of the on-treatment count, although the two are not independent and the subgroups are small.

**Figure 5:**
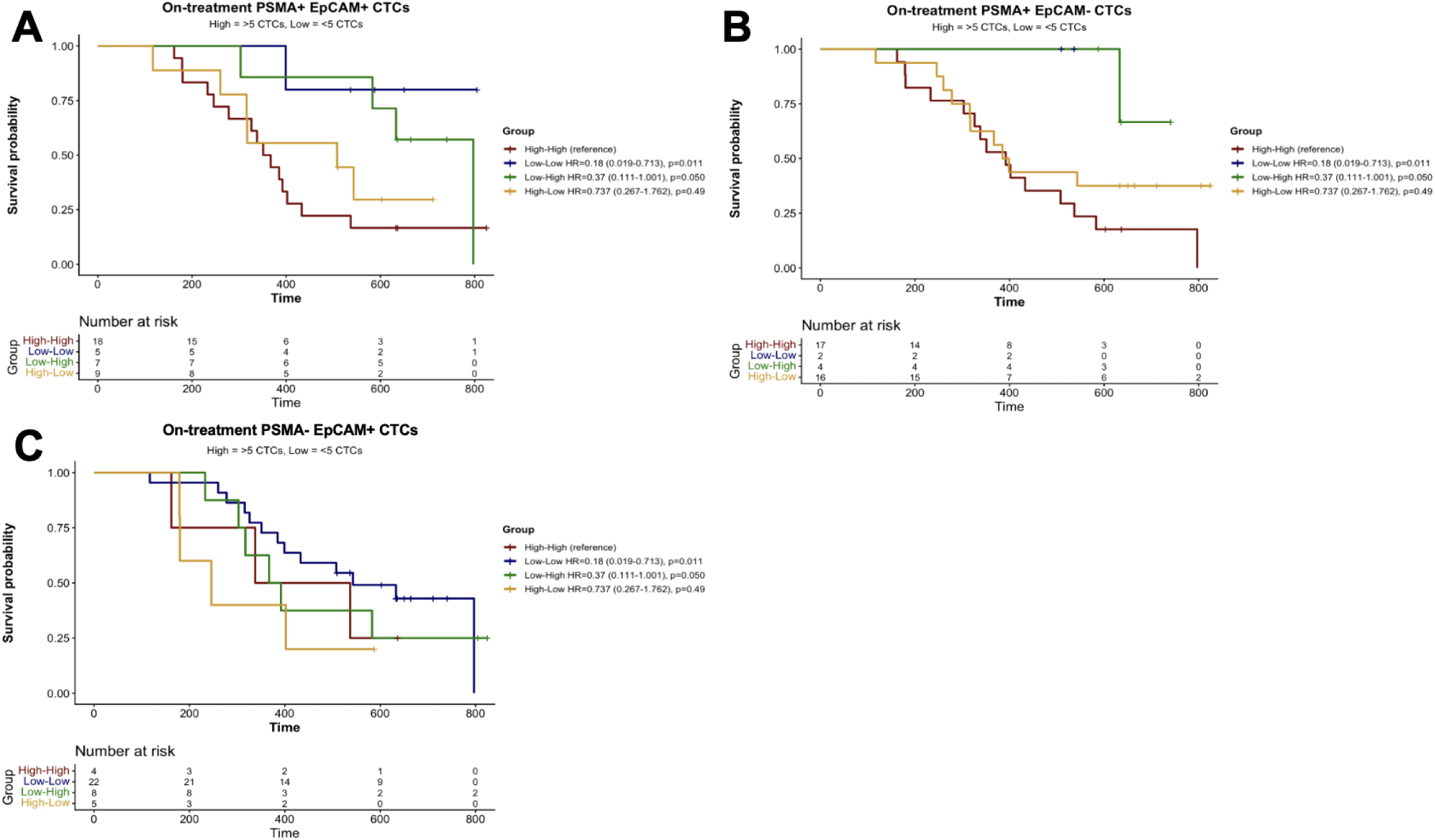
Changes in CTC subset counts between pre- and on-treatment samples are associated with survival outcomes. Kaplan-Meier survival curves showing overall survival based on stratification by change in the number of CTCs recovered between pre- and on-treatment samples. Patients were divided into four groups: High-High (>5 CTCs in both pre- and on-treatment samples); Low-Low (<=5 CTCs in both pre- and on-treatment samples); Low-High (<=5 CTCs in pre-treatment sample and >5 CTCs in on-treatment sample); and High-Low (>5 CTCs in pre-treatment sample and <=5 CTCs in on-treatment sample). Data are shown for PSMA+ EpCAM+ CTCs (A), PSMA+ EpCAM-CTCs (B), and PSMA-EpCAM+ CTCs (C).

The only condition shown to have an unfavorable hazard ratio relative to the High-High group was the High-Low group within the PSMA−/EpCAM+ subset of CTCs (HR=1.53, 95%CI 0.37-6.91, p=0.55), although having a lower number of PSMA-EpCAM+ CTCs at the on-treatment time point was indicative of longer overall survival (Fig. 5C). The Low-Low group for this subset had a more favorable hazard ratio relative to the High-High group (HR=0.55, 95%CI 0.19-2.12, p=0.348) and relative to the High-Low group (HR=0.36, 95%CI 0.13-1.19, p=0.089), suggesting that low PSMA-CTC counts at the on-treatment time point are not themselves associated with worse overall survival; rather, a decrease in this population may correlate with less favorable outcomes. None of these comparisons reached statistical significance, and each subgroup contained fewer than ten patients, so this observation should be interpreted as hypothesis-generating.

## Discussion

Here, we show that distinct subsets of CTCs determined by presence of PSMA and EpCAM exist in mCRPC patients and we demonstrate that these populations are differentially affected by ^177^Lu-PSMA-617 treatment, carrying implications for future prognostic tools and the development of post-^177^Lu-PSMA-617 therapies.

To our knowledge, this is the first study to evaluate CTC subsets defined by PSMA and EpCAM as prognostic indicators in patients with mCRPC receiving ^177^Lu-PSMA-617. Analysis of these subsets was associated with overall survival, with lower numbers of PSMA+ CTCs in on-treatment samples being associated with more favorable overall survival outcomes.

The data herein also constitute, to our knowledge, the first analysis to correlate cell-surface PSMA expression dynamics on tumor cells with survival in the setting of ^177^Lu-PSMA-617 treatment and only the second to directly assess protein-level PSMA on CTCs in this context^24^. ^177^Lu-PSMA-617’s mechanism of action centers on the irradiation of PSMA+ tumor cells^23^; therefore, it is unsurprising that we observed a reduction in the proportion and abundance of PSMA+ cells between the pre- and on-treatment samples in this study. A greater fraction of PSA50 responders (15/19) than non-responders (13/20) had a numerical decrease in the total number of PSMA+ CTCs recovered between their pre- and on-treatment samples. This finding is consistent with a recent study that found that both PSA responders and non-responders experienced a decline in PSMA MFI over the course of three cycles of ^177^Lu-PSMA-617^24^. However, our analyses reveal several notable nuances. First, while there were numerical decreases in both PSMA+/EpCAM+ CTCs and PSMA+/EpCAM-CTCs, the proportion of PSMA+/EpCAM-CTCs decreased while the fraction of PSMA+/EpCAM+ cells slightly increased between the pre- and on-treatment time points. This may indicate that EpCAM-tumor cells are more susceptible to ^177^Lu-PSMA-617 than EpCAM+ cells, effectively enriching the tumor population for EpCAM+ cells, although this interpretation is speculative and was not directly tested here. EpCAM expression has been shown to correlate with higher Gleason score in prostate cancer^25^ and increased abundance of EpCAM+ CTCs is correlated with worse survival outcomes in mCRPC^26,27^. However, EpCAM is also typically lost during epithelial to mesenchymal transition as prostate cancer progresses^28^. Prior studies have often defined CTCs as EpCAM+ cells^13,19,29,30^. The platform employed in our study is advantageous in this context because it enables simultaneous detection of multiple surface proteins on CTCs without relying on markers such as EpCAM for identification or the need for additional downstream analyses^20^.

Next, we demonstrate that PSMA+ tumor cells remain in circulation following ^177^Lu-PSMA-617 treatment, even in PSA50 responders. This is highly relevant to the development of post-^177^Lu-PSMA-617 therapies, as it is currently debated whether patients who have responded to this treatment would subsequently be susceptible to other PSMA-targeting strategies such as other PSMA-directed radioligands^31^, T cell engagers^32^, and CAR T cells^33^. Our results suggest that a population of PSMA+ cells remain in circulation following multiple cycles of ^177^Lu-PSMA-617, although additional studies should assess the proportion and number of PSMA+ CTCs after a full course of treatment is complete.

Finally, our study identified a previously undescribed increase in the proportion and absolute count of PSMA−/EpCAM+ CTCs following ^177^Lu-PSMA-617 treatment. This likely reflects the selective killing of PSMA+ cells by the PSMA-targeting radioligand therapy and the subsequent reciprocal increase in PSMA-tumor cells. Interestingly, while patients with a lower number of these PSMA-CTCs at the on-treatment time point still had favorable survival outcomes relative to those with higher numbers, this trend was much weaker in the PSMA-population than in either PSMA+ subset. Furthermore, when we examined the effects of changes in each CTC population on survival, the PSMA-compartment was the only subset in which patients who had sustained high levels of those CTCs did not have the worst survival outcomes; instead, patients who experienced a decrease in PSMA-cells between pre- and on-treatment time points had the shortest OS of any group. This comparison was not statistically significant (p=0.55) and involved a small number of patients, so it should be interpreted with caution.

Another recent study examining the transcriptomic landscape of CTCs in mCRPC also included a subset of patients who were prospectively enrolled in a longitudinal study of CTCs in the setting of ^177^Lu-PSMA-617 therapy^13^. Importantly, CTCs in this study were defined as EpCAM+. The authors found that CTCs with a luminal B subtype-like transcriptomic profile in pre-treatment samples were associated with unfavorable survival outcomes and a lower clinical response rate to ^177^Lu-PSMA-617. The expression of PSMA on this population was similar to that of other subsets that had more favorable response rates, providing a possible explanation for the observation that the dynamics of PSMA+ EpCAM+ CTCs were similar between PSA50 responders and non-responders in our study. Future studies should seek to explore this further by assessing the distinct transcriptomic profiles of PSMA+/EpCAM+, PSMA+/EpCAM-, and PSMA−/EpCAM+ CTCs in mCRPC.

## Conclusions

Collectively, the results of this study support the further evaluation of PSMA and EpCAM expression on CTCs as a potential prognostic indicator during ^177^Lu-PSMA-617 treatment, and provide an early characterization of PSMA and EpCAM expression dynamics on CTCs in this setting. Prospective validation in larger, multi-institutional cohorts, ideally with sampling at the completion of therapy and at progression, is warranted.

## Study limitations

This study has several limitations that should be considered when interpreting its results. First, this analysis was performed on a small cohort of only 39 patients with available serial CTC samples and was conducted solely at one institution, potentially restricting its generalizability. Future studies should seek to validate these results in additional patients and at other sites. Second, the on-treatment samples analyzed were collected partway through ^177^Lu-PSMA-617 treatment (between 12 and 24 weeks after initiation) rather than at the end of the therapeutic process, limiting the scope of its conclusions relating to the landscape of tumor cell PSMA protein at the time of treatment failure or disease progression. Finally, CTCs were stratified as PSMA+/- and EpCAM+/- in a binary fashion, precluding the assessment of the magnitude of surface levels and any analysis of the association between protein levels and survival or biochemical response outcomes. Further biological information could also have been captured by interrogating tumor-cell DNA and RNA (including analysis of signatures or pathways), which was beyond the scope of this study. In addition, the survival analyses were univariable and were not adjusted for established prognostic factors such as baseline PSA, alkaline phosphatase, ECOG performance status, or number of prior lines of therapy; the number of deaths was small relative to the number of comparisons performed; and the PSA50 responder and non-responder groups were defined post hoc. The survival findings should therefore be regarded as hypothesis-generating rather than confirmatory.

## Conflict of Interest Statement

J.M.D. serves as Chief Scientific Officer of Astrin Biosciences, the company that developed the circulating tumor cell detection platform used in this study, and holds stock options in that company. This interest has been reviewed and managed by the University of Minnesota in accordance with its Conflict-of-Interest policies. M.J.L., L.Y., S.D., J.H.H., and A.T.A. declare no competing interests relevant to this work. N.Z. Honoraria: Association of Community Cancer Centers (ACCC)/MJH, Curio Science, Department of Defense Prostate Cancer Research Program, Guidepoint Global, MJH, Mosaic Research Management, Slingshot Insights, Bayer (Institutional), Vaniam Group; Advisory: Bayer (Personal); Research Funding (to Institution): Janssen Research & Development, Johnson & Johnson, Janux Therapeutics, Amgen, Lava Therapeutics, GT Biopharma, ArsenalBio, Takeda; Travel, Accommodations and Expenses (Expense only): Telix Pharmaceuticals, Caris Life Sciences, Bayer, Janux Therapeutics, DAVA Oncology; Uncompensated Relationships: Takeda, Amgen (Advisory). E.S.A. reports grants and personal fees from Janssen, Johnson & Johnson, Sanofi, Bayer, Bristol Myers Squibb, Convergent Therapeutics, Curium, MacroGenics, Merck, Pfizer, and AstraZeneca; personal fees from Aadi Bioscience, Abeona Therapeutics, Aikido Pharma, Astellas, Amgen, Blue Earth, Boundless Bio, Corcept Therapeutics, Duality Bio, Exact Sciences, Hookipa Pharma, Invitae, Eli Lilly, Foundation Medicine, Menarini-Silicon Biosystems, Tango Therapeutics, Tempus, Tolmar Scientific, VIR Biotechnology, and Z-alpha; grants from Novartis, Celgene, and Orion; and has a patent for an AR-V7 biomarker technology that has been licensed to Qiagen. M.J.L. has no conflicts to disclose.

---

**Supplemental Table 1:**
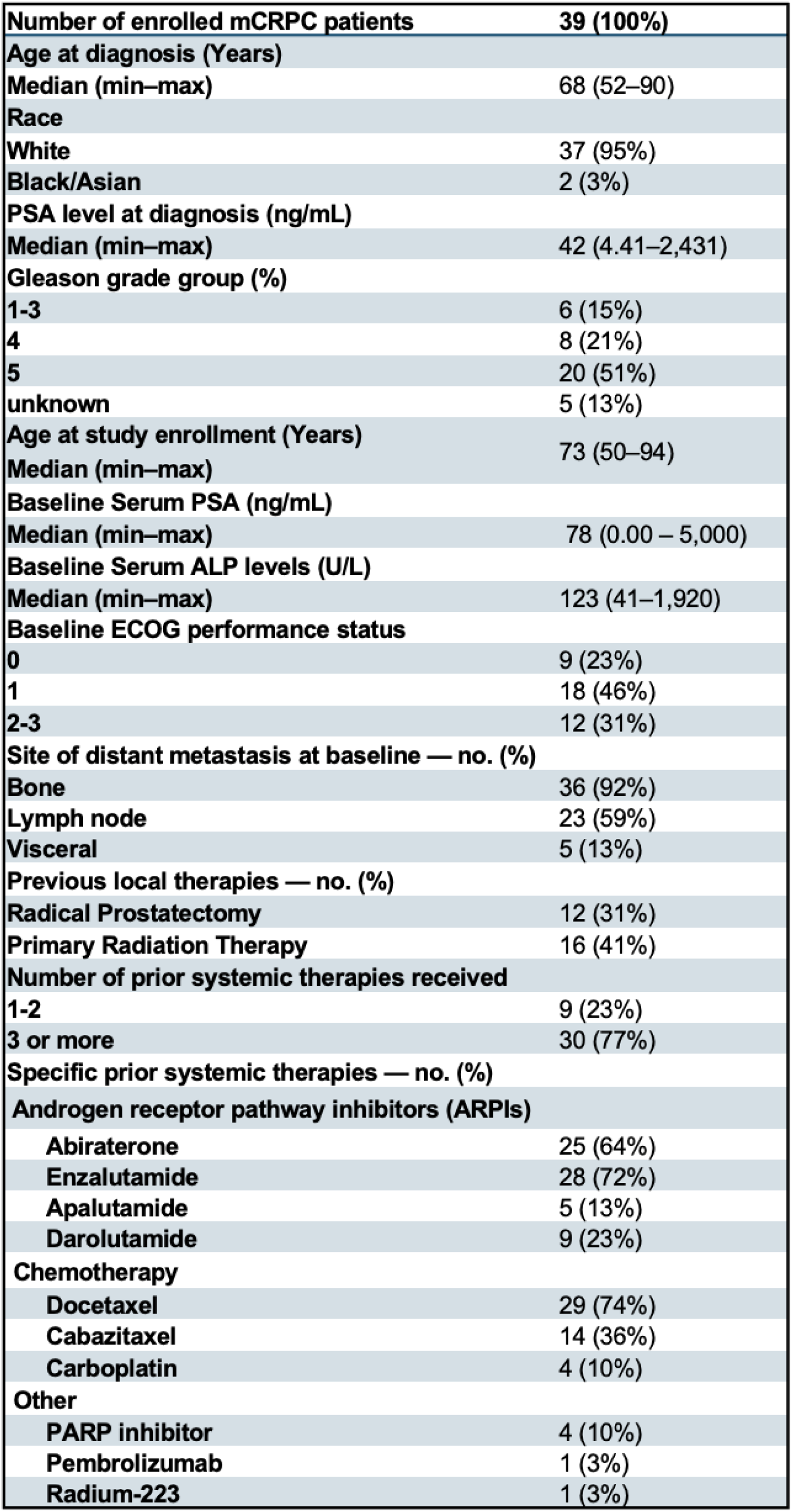
Baseline characteristics of mCRPC patients enrolled in this study.

