## Supplementary figures and images for "Dynamics of circulating tumor cell subsets defined by PSMA and EpCAM predict survival in metastatic castration-resistant prostate cancer patients treated with ^177^Lu-PSMA-617"

### Supplemental Figure 1

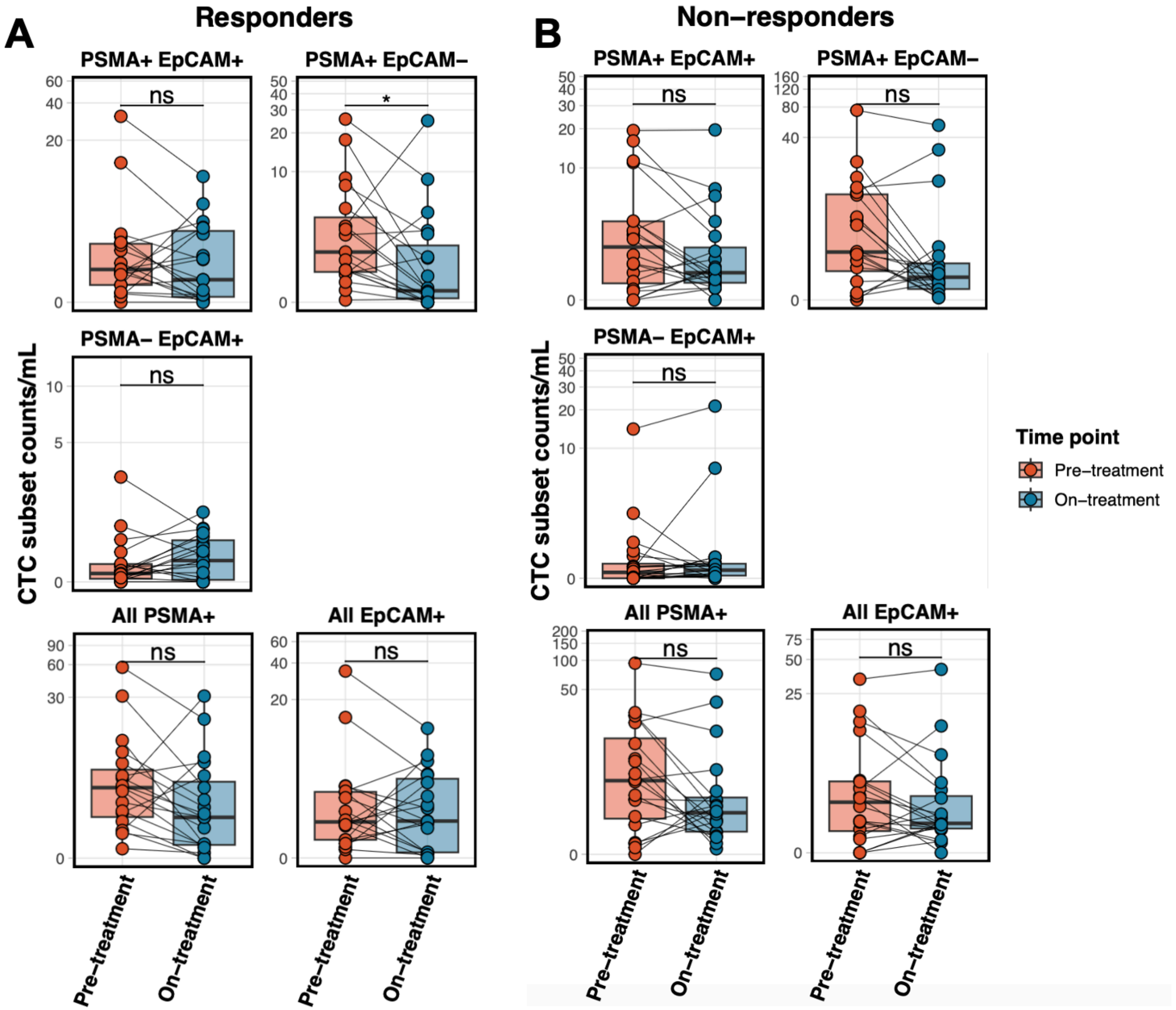
