## Supplemental Table 1 for "Dynamics of circulating tumor cell subsets defined by PSMA and EpCAM predict survival in metastatic castration-resistant prostate cancer patients treated with ^177^Lu-PSMA-617"

| **Number of enrolled mCRPC patients** | **39 (100%)** |
| --- | --- |
| **Age at diagnosis (Years)** |  |
| **Median (min–max)** | 68 (52–90) |
| **Race** |  |
| **White** | 37 (95%) |
| **Black/Asian** | 2 (3%) |
| **PSA level at diagnosis (ng/mL)** |  |
| **Median (min–max)** | 42 (4.41–2,431) |
| **Gleason grade group (%)** |  |
| **1-3** | 6 (15%) |
| **4** | 8 (21%) |
| **5** | 20 (51%) |
| **unknown** | 5 (13%) |
| **Age at study enrollment (Years)**  **Median (min–max)** | 73 (50–94) |
| **Baseline Serum PSA (ng/mL)** |  |
| **Median (min–max)** | 78 (0.00 – 5,000) |
| **Baseline Serum ALP levels (U/L)** |  |
| **Median (min–max)** | 123 (41–1,920) |
| **Baseline ECOG performance status** |  |
| **0** | 9 (23%) |
| **1** | 18 (46%) |
| **2-3** | 12 (31%) |
| **Site of distant metastasis at baseline — no. (%)** |  |
| **Bone** | 36 (92%) |
| **Lymph node** | 23 (59%) |
| **Visceral** | 5 (13%) |
| **Previous local therapies — no. (%)** |  |
| **Radical Prostatectomy** | 12 (31%) |
| **Primary Radiation Therapy** | 16 (41%) |
| **Number of prior systemic therapies received** |  |
| **1-2** | 9 (23%) |
| **3 or more** | 30 (77%) |
| **Specific prior systemic therapies — no. (%)** |  |
| **Androgen receptor pathway inhibitors (ARPIs)** |  |
| **Abiraterone** | 25 (64%) |
| **Enzalutamide** | 28 (72%) |
| **Apalutamide** | 5 (13%) |
| **Darolutamide** | 9 (23%) |
| **Chemotherapy** |  |
| **Docetaxel** | 29 (74%) |
| **Cabazitaxel** | 14 (36%) |
| **Carboplatin** | 4 (10%) |
| **Other** |  |
| **PARP inhibitor** | 4 (10%) |
| **Pembrolizumab** | 1 (3%) |
| **Radium-223** | 1 (3%) |
